# Rapid and precise genome engineering in a naturally short-lived vertebrate

**DOI:** 10.1101/2022.05.25.493454

**Authors:** Ravi D. Nath, Claire N. Bedbrook, Rahul Nagvekar, Karl Deisseroth, Anne Brunet

## Abstract

The African turquoise killifish is a powerful vertebrate system to study complex phenotypes at scale, including aging and age-related disease. Here we develop a rapid and precise CRISPR/Cas9-mediated knock-in approach in the killifish. We show its efficient application to precisely insert fluorescent reporters of different sizes at various genomic loci, to drive cell-type- and tissue-specific expression. This knock-in method should allow the establishment of humanized disease models and the development of cell-type-specific molecular probes for studying complex vertebrate biology.

## Main text

Studying complex biological phenotypes such as aging and disease in vertebrates is limited by issues of scale and speed. For example, the inherent long lifespan and low-throughput nature of mice prohibit iterative genetics and exploration of vertebrate biology. The African turquoise killifish *Nothobranchius furzeri* (hereafter killifish) has emerged as a powerful model to overcome this challenge and accelerate discovery due to its rapid timeline for sexual maturity (3–4 weeks post hatching) and naturally compressed lifespan (4–6 months) (Hu and Brunet, 2018; Kim et al., 2016). The killifish has the shortest generation time of a vertebrate model system bred in the laboratory (2 months) (Hu and Brunet, 2018; Kim et al., 2016; Polacik et al., 2016), making rapid vertebrate genetics possible. Tools to advance genetic interrogation of the killifish have been developed, including a sequenced genome (Reichwald et al., 2015; Valenzano et al., 2015) and Tol2 transgenesis (Allard et al., 2013; Hartmann and Englert, 2012; Valenzano et al., 2011), as well as CRISPR/Cas9-mediated knock-out (Harel et al., 2015) and CRISPR/Cas13-mediated knock-down (Kushawah et al., 2020). This genetic toolkit has enabled discoveries about the mechanisms of aging (Astre et al., 2022a; Bradshaw et al., 2022; Chen et al., 2022; Harel et al., 2022; Louka et al., 2022; Matsui et al., 2019; Smith et al., 2017; Van Houcke et al., 2021b), regeneration (Vanhunsel et al., 2022a; Vanhunsel et al., 2021; Vanhunsel et al., 2022b; Wang et al., 2020), evolution (Cui et al., 2019; Sahm et al., 2017; Singh et al., 2021; Willemsen et al., 2020), development (Abitua et al., 2021; Dolfi et al., 2019), and ‘suspended animation’ (Hu et al., 2020; Singh et al., 2021).

Knock-in methods are essential for genetic tractability of model organisms. They enable precise mutations in key genes for mechanistic studies and human disease modeling. Knock-in technologies also allow the insertion of molecular tags or reporters at specific genomic loci. Combined with self-cleaving peptides, a knock-in approach can be leveraged to drive cell-type-and tissue-specific expression of ectopic genes (e.g., genes of interest, recombinases) or probes (e.g., fluorescent reporters, calcium indicators). While small insertions (<8 bp) have been achieved via knock-in in the killifish genome in the parental generation (Harel et al., 2015), a method to precisely insert large transgenes and allow the efficient generation of stable lines with germline transmission is missing.

### CRISPR/Cas9-mediated knock-in in killifish allows efficient tissue-specific expression of fluorescent reporters

To achieve precise integration of genes of interest at endogenous target loci, we developed a method based on CRISPR/Cas9-mediated homology-directed repair (HDR). CRISPR/Cas9-mediated HDR is often associated with issues of low efficiency and multicopy insertion (Auer et al., 2014). To overcome these issues, we injected killifish embryos with a cocktail (see Methods) composed of (1) recombinant Cas9 protein, (2) chemically-modified guide RNAs (gRNAs), (3) a chemically-modified linear double-stranded DNA (dsDNA) HDR template, and (4) an HDR chemical enhancer which inhibits non-homologous end joining (NHEJ) (DiNapoli et al., 2020; Gutierrez-Triana et al., 2018; Seleit et al., 2021; Wierson et al., 2020). We designed the dsDNA HDR template with 150–200 bp homology arms flanking the site of insertion at the target locus (in this case the stop codon of a specific gene) (Figure 1A). To rapidly assess the efficiency of CRISPR/Cas9-mediated knock-in, we included the following sequences in the dsDNA HDR template: a *T2A* sequence (encoding the T2A self-cleaving peptide (Szymczak et al., 2004)) and the fluorescent protein Venus (Nagai et al., 2002). Use of the T2A self-cleaving peptide avoids direct fusion of the fluorescent protein to the targeted gene’s protein product (Figure 1A; Supplemental Table 1). The modified dsDNA HDR template and gRNAs can all be directly ordered (see Methods), which alleviates the need for cloning or PCR. With successful insertion, the expression of Venus should be controlled by the endogenous regulatory elements (e.g., promoter, enhancers) of the target gene, which could be leveraged for cell-type-or tissue-specific expression.

**Figure 1:**
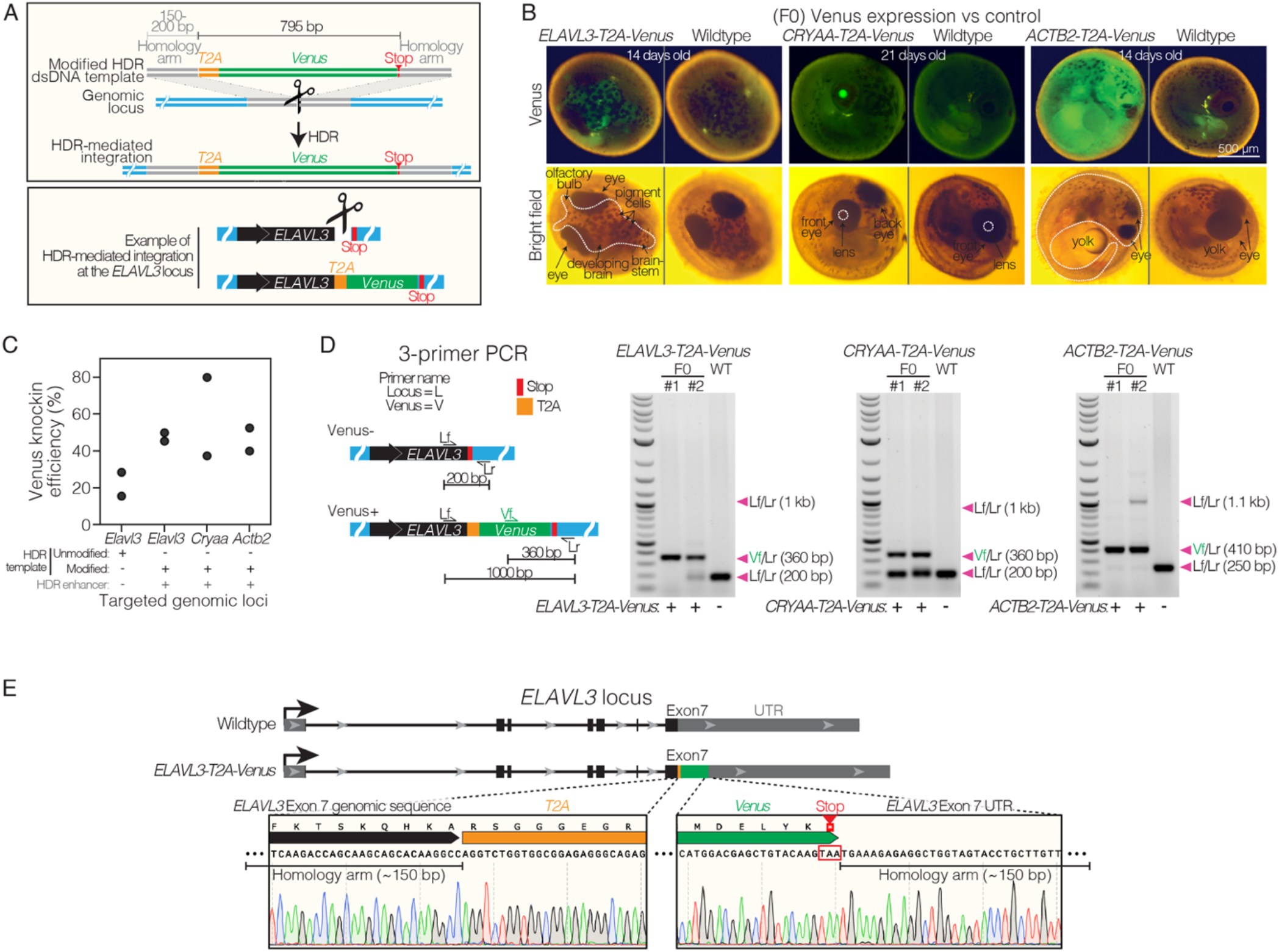
Efficient homology directed repair for precise knock-in at different genomic locations in killifish. A. Schematic of *T2A*-*Venus* insertion at the *ELAVL3* locus. B. Images of F0 Venus+ and wildtype 14–21-day-old embryos for each targeted locus (*ELAVL3, CRYAA*, and *ACTB2*). Twenty-one-day-old embryos were dried on coconut fiber for 7 days prior to imaging and have altered autofluorescence compared to 14-day-old embryos that were not yet put on coconut fiber. C. Efficiency of *T2A*-*Venus* knock-in at each locus (determined by visual inspection of Venus fluorescence in developed embryos) and efficiency of knock-in at *ELAVL3* with a dsDNA HDR template lacking chemical modification and without the HDR chemical enhancer; two replicates per condition; *n* = 10–42 embryos per replicate. Raw data in Supplemental Table 2. D. Left, 3-primer PCR schematic showing locus-specific external primers forward (Lf) and reverse (Lr) and internal forward Venus primer (Vf). Right, gel images of 3-primer PCR for each locus comparing F0 with wildtype (WT) fish. Arrowheads indicate each primer pair and its expected amplification product length. Scoring Venus positive (+) or negative (-) for each fish is indicated below the gel images. Note that the relatively large ∼1 kb Lf/Lr product in the transgenic F0 animals is likely to be outcompeted by the shorter Vf/Lr amplification product during the PCR reaction. E. Top, comparison of *ELAVL3* locus for wildtype and *ELAVL3-T2A-Venus*. Bottom, precise in-frame insertion of *T2A-Venus* in exon 7, immediately before the stop codon of *ELAVL3* and followed by the *ELAVL3* untranslated region (UTR).

Using this approach, we targeted Venus to three distinct genomic loci in the killifish: *ELAVL3, CRYAA*, and *ACTB2*, which are known to have brain-specific (Ahrens et al., 2012), lens-specific (Posner et al., 2017), and ubiquitous (Gutierrez-Triana et al., 2018) expression, respectively, in teleost fish (including zebrafish and medaka). After injection of CRISPR/Cas9 reagents into one-cell stage killifish embryos, we waited 14–21 days for the embryos to develop and visually screened embryos for Venus fluorescence – indicative of protein expression and suggestive of successful CRISPR/Cas9-mediated knock-in. We observed Venus fluorescent protein expression in the expected tissues: developing brain for *ELAVL3-*targeted embryos, lens of the eye for *CRYAA-*targeted embryos, and in all cells of the embryo for *ACTB2*-targeted embryos (Figure 1B). In all embryos screened, we did not observe Venus expression in a tissue that was not specifically targeted. For all three targeted loci, we observed Venus fluorescence (suggestive of successful CRISPR/Cas9-mediated knock-in) in over 40% of developed embryos (Figure 1C). We achieved the highest CRISPR/Cas9-mediated knock-in efficiency using both a chemically-modified dsDNA HDR template (modification #3, see Methods) and HDR chemical enhancer compared to the use of unmodified HDR template without enhancer for the *ELAVL3* locus (Figure 1C; Figure 1 — figure supplement 1A), so we used this approach for all subsequent constructs. We did not observe differences in lethality of embryos injected with CRISPR/Cas9 knock-in reagents (including chemically-modified dsDNA HDR template and/or HDR chemical enhancer) compared to non-injected embryos (Figure 1—figure supplement 1B). Importantly, we confirmed that the genomic knock-in occurred at the expected genomic locus by PCR genotyping with primers surrounding the insertion site for each gene (Figure 1D; Supplemental Table 1) (see below for sequencing confirmation in the F1 generation). Thus, this CRISPR/Cas9-mediated knock-in method allows for precise and efficient editing at several loci, including tissue-specific ones.

### Germline transmission of CRISPR/Cas9-mediated knock-in and generation of stable lines

A key aspect of genome editing is germline transmission to allow the generation of genetically-modified lines. To determine if the CRISPR/Cas9-mediated insertion can be transmitted to the next generation, we evaluated the efficiency of germline transmission using transgenic *ELAVL3*-*T2A*-*Venus* founders (F0s). Sixty-seven percent of F0 founders, when crossed with wildtype fish, produced Venus-positive F1 progeny (Figure 2A, B). Given the high efficiency of germline transmission and the rapid generation time of killifish, we tested if we could directly generate homozygous F1 animals by inter-crossing genetically modified F0 individuals (Figure 2C, D). Upon inter-crossing Venus-positive F0 founders, we found that 85% of the resulting F1 Venus-positive progeny were homozygous for the insertion at the *ELAVL3* locus (Figure 2D). PCR amplification and genotyping by Sanger sequencing of homozygous F1 animals confirmed that the *T2A*-*Venus* integration at the *ELAVL3* locus was as designed—single-copy and in frame (Figure 1E; Figure 2A, C, D; Figure 2—figure supplement 1). Venus-positive *ELAVL3*-*T2A*-*Venus* F1 progeny exhibit specific and strong Venus expression throughout the nervous system including the retina, brain, and spinal cord at the larval stage (Figure 2E), which is expected given that the *ELAVL3* promoter is commonly used as a pan-neuronal promoter in larval zebrafish (Ahrens et al., 2012) (see Figure 3 below for expression in adult brains). We tested for potential off-target insertions/mutations upon CRISPR/Cas9-mediated knock-in. PCR amplification and Sanger sequencing of homozygous F1 *ELAVL3*-*T2A*-*Venus* animals at the three most likely off-target sites (predicted by CHOPCHOP (Labun et al., 2019)) showed no off-target editing at these sites (Figure 2— figure supplement 2). While there could be insertions at other sites in the genome, the observation that the expression pattern of Venus recapitulates that of the known endogenous gene (for *ELAVL3, CRYAA, ACTB2*, and other loci, see Figure 4) supports the notion that in-frame off-target insertions are rare with this method. In rare cases where off-target insertions would occur, they could be eliminated by backcrossing lines with wildtype fish. Hence, this method enables generation of stable lines of homozygous transgenic vertebrate animals in 2–3 months.

**Figure 2:**
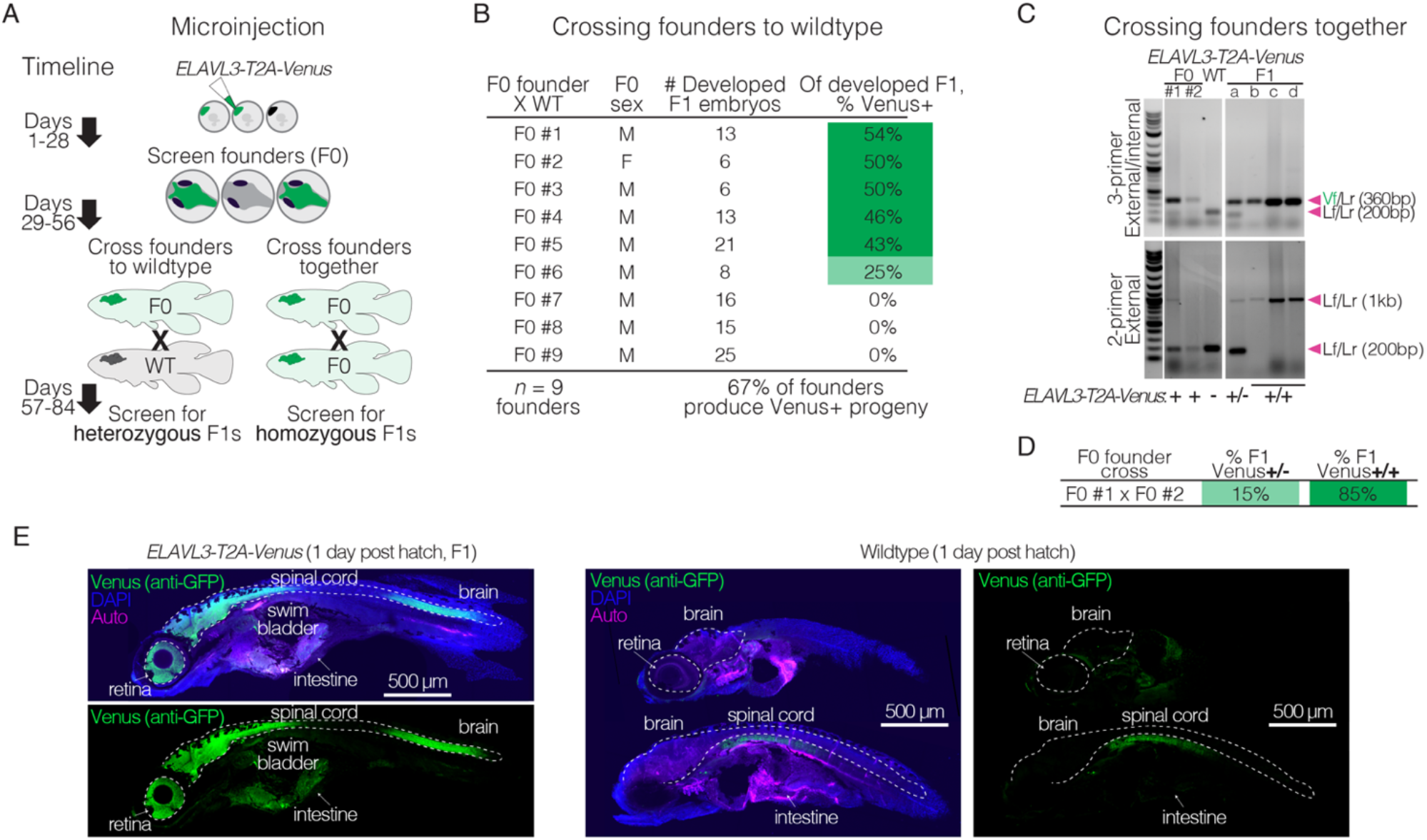
Rapid generation of stable knock-in lines in the killifish. A. Schematic of generating a stable knock-in line by either crossing F0 X WT (left) or F0 X F0 (right), with timelines to verified heterozygous or homozygous animals. B. Germline transmission of *T2A*-*Venus* at the *ELAVL3* locus was verified by crossing F0 X WT (*n* = 9 breeding pairs), with between 6–25 developed embryos per pair screened visually or by PCR. C. Crossing F0 animals positive for *T2A*-*Venus* at the *ELAVL3* locus (F0 X F0). Gels showing F0 parents (left) and F1 progeny (a, b, c, and d; right) with 3-primer PCR (top) and external PCR (bottom) using Venus and locus-specific primers shown in Figure 1D. The external PCR shows both heterozygous (a) and homozygous (b, c, and d) F1 progeny for *ELAVL3-T2A*-*Venus*. Arrowheads indicate each primer pair and its expected amplification product length. Scoring for each lane of the gel is indicated below the gel images. F0 animals are likely mosaic so only “+” or “-” was assigned based on the 3-primer PCR result. D. Percent of fully developed and Venus+ F1 progeny from the F0 X F0 cross that are heterozygous (+/-) or homozygous (+/+) for insertion of *Venus* in *ELAVL3-T2A-Venus* animals. E. Sagittal sections of F1 homozygous *ELAVL3-T2A*-*Venus* (left) compared with wildtype (right) killifish 1 day post hatch (larval stage) showing merge of Venus (stained with anti-GFP antibody; green), DAPI (nuclei; blue), and autofluorescence (‘Auto’; magenta) as well as separate channel for Venus (stained with anti-GFP antibody; green). Scale bar = 500 μm.

**Figure 3:**
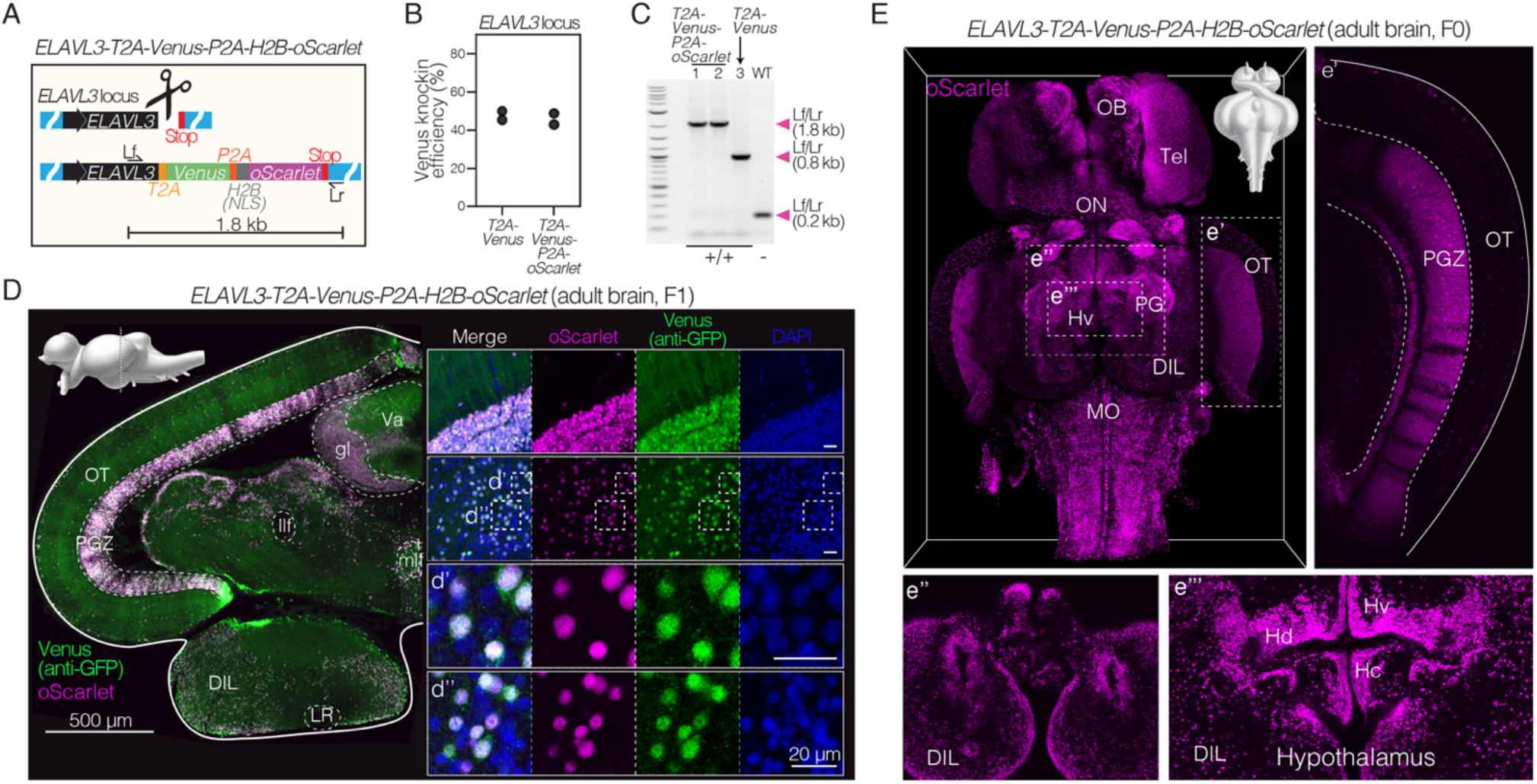
Efficient and stable knock-in of a large 1.8 kb insertion in killifish. A. Schematic of design of a *T2A*-*Venus*-*P2A*-*H2B*-*oScarlet* sequence for targeted knock-in at the *ELAVL3* locus and locus-specific external primers forward (Lf) and reverse (Lr). B. Knock-in efficiency comparing 1.8 kb insertion (*ELAVL3*-*T2A*-*Venus*-*P2A*-*H2B*-*oScarlet*) to the 0.8 kb insertion (*ELAVL3*-*T2A*-*Venus*) determined by visual inspection of developed embryos for Venus fluorescence; two independent replicates per condition; *n* = 10–86 embryos per replicate. Raw data in Supplemental Table 2. C. PCR amplification at the *ELAVL3* locus using locus-specific external primers forward (Lf) and reverse (Lr) shown in (A) comparing amplicon length from two F1 *ELAVL3*-*T2A*-*Venus*-*P2A*-*H2B-oScarlet* animals (lane 1 and 2), one F1 *ELAVL3*-*T2A*-*Venus* animal (lane 3), and one wildtype animal (lane WT), showing a single band at the expected length in each case. Scoring for each lane of the gel is indicated below the gel image. D. Left, coronal brain section of adult (3 months old) *ELAVL3*-*T2A*-*Venus*-*P2A*-*H2B-oScarlet* heterozygous F1 male, showing expression of Venus and oScarlet. Scale bar = 500 μm. Upper left corner, sagittal view of the *N. furzeri* brain adapted from (D’Angelo, 2013) indicating the plane of the coronal section. Right, select regions showing separate channels for oScarlet (magenta), Venus (stained with anti-GFP antibody; green), DAPI (nuclei; blue) as well as merged channels. (d’) and (d’’): zoomed in individual cells. Scale bar = 20 μm. oScarlet expression is confined to nuclei while Venus expression is observed throughout cell bodies and projections. E. Brain-wide expression of nuclear-localized oScarlet (magenta) in adult (1 month old) *ELAVL3*-*T2A*-*Venus*-*P2A*-*H2B-oScarlet* F0 male. Select regions are highlighted: (e’) the optic tectum (OT), (e’’) the most ventral view of the hypothalamus, and (e’’’) the periventricular hypothalamus.

**Figure 4:**
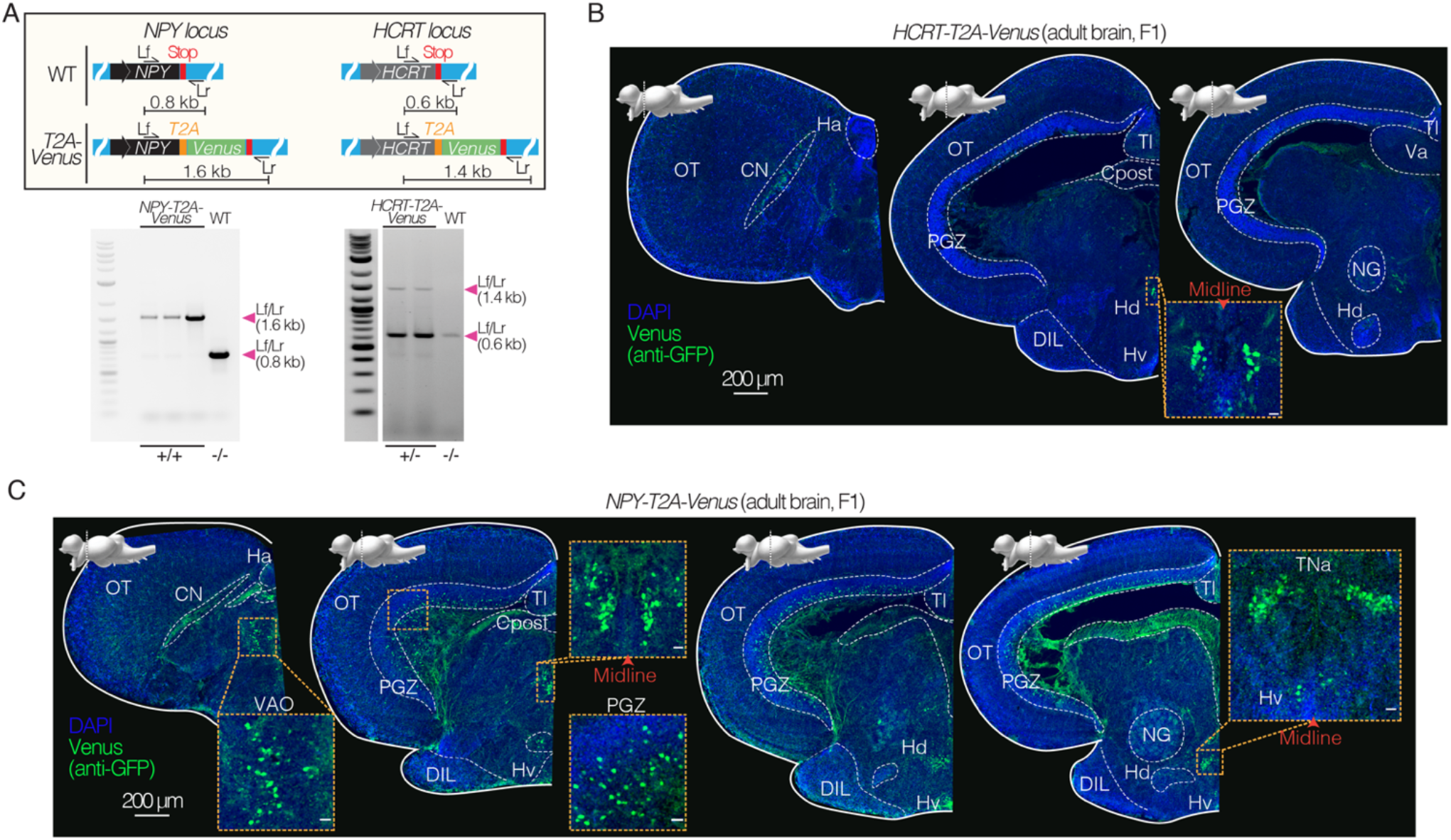
Expression in specific neuronal populations using CRISPR/Cas9 knock-in lines in killifish. A. Top, schematics of design of *T2A*-*Venus* sequence for targeted knock-in at the *NPY* and *HCRT* loci including locus-specific external primers forward (Lf) and reverse (Lr). Bottom, PCR amplification at the *NPY* or *HCRT* locus comparing amplicon length from *NPY*-*T2A*-*Venus* (F1 animals) versus wildtype (WT) and comparing amplicon length from *HCRT*-*T2A*-*Venus* (F1 animals) versus wildtype (WT). B. Coronal brain sections of adult (4 months old) *HCRT*-*T2A*-*Venus* female (heterozygous F1), showing Venus expression (stained with anti-GFP antibody; green) and DAPI (nuclei; blue). Scale bar = 200 μm. Distinct nuclei indicated and labeled with abbreviated names. Above each slice is the sagittal view of the *N. furzeri* brain adapted from (D’Angelo, 2013) indicating the plane of the coronal section. Inset shows zoom in on Venus positive population of cells in the dorsal hypothalamus close to the midline. Scale bar = 20 μm. C. Coronal brain sections of adult (3.5 months old) *NPY*-*T2A*-*Venus* male (homozygous F1), showing Venus expression (stained with anti-GFP antibody; green) and DAPI (nuclei; blue). Scale bar = 200 μm. Distinct nuclei indicated and labeled with abbreviated names. Above each slice is the lateral view of the *N. furzeri* brain adapted from (D’Angelo, 2013) indicating the plane of the coronal section. Insets show zoom in on the Venus positive populations. Scale bar = 20 μm.

### Insertion of long sequences into the genome to drive gene expression in a cell-or tissue-specific manner

We asked if this CRISPR/Cas9-mediated insertion method could be used to insert longer sequences into the killifish genome for expression in specific cells or tissues. The insertion of long sequences at a precise genomic location, while technically challenging, is critical for leveraging the cell-type or tissue specificity of a particular locus to drive ectopic expression of specific genes or molecular probes. We designed a longer dsDNA HDR template that would result in a 1.8 kb long insertion sequence. This HDR template includes two consecutive fluorescent proteins (Venus and oScarlet) targeted to the *ELAVL3* locus, with *T2A* and *P2A* sequences (encoding another self-cleaving peptide) 5’ to each fluorescent protein, respectively, to avoid direct fusion. The oScarlet was also tagged with the nuclear localization signal (NLS) from Histone 2B to allow nuclear localization of this fluorescent protein (Kanda et al., 1998; Schrodel et al., 2013). The resulting insertion sequence is 1.8 kb long – a length that would encode proteins of ∼600 amino acids and ∼65 kDa (Figure 3A; Supplemental Table 1). We observed successful CRISPR/Cas9-mediated knock-in of this longer sequence in ∼50% of developed embryos (Figure 3B). There was no decrease in efficiency for this longer insertion relative to the shorter (0.8 kb) insertion previously tested at the same locus (Figure 3B). PCR amplification and genotyping by Sanger sequencing of homozygous F1 animals confirmed that the *T2A*-*Venus-P2A-H2B-oScarlet* integration at the *ELAVL3* locus was the expected size and in frame without mutations (Figure 3C; Supplemental Table 1). PCR amplification and Sanger sequencing of homozygous F1 animals at the three predicted most likely off-target sites showed no off-target editing in these fish either (Figure 3—figure supplement 1). Imaging coronal brain sections of adult F1 *ELAVL3-T2A*-*Venus-P2A-H2B-oScarlet* killifish showed cells (likely neurons) expressing both oScarlet and Venus (Figure 3D). As expected, oScarlet expression was confined to nuclei while Venus expression was seen in both cell bodies and projections (Figure 3D). Imaging the whole brain of adult *ELAVL3*-*T2A*-*Venus*-*P2A*-*H2B*-*oScarlet* killifish revealed oScarlet-positive nuclei throughout the brain (Figure 3E). Thus, this method allows for pan-neuronal expression in the adult brain and could be leveraged to drive expression of molecular tools (e.g., the optogenetic ion channel channelrhodopsin [∼1 kb] (Boyden et al., 2005) or the genetically-encoded calcium indicator GCaMP [∼1.3 kb] (Ahrens et al., 2012)) in a neuronal-specific manner.

### Cell-type-specific expression in subsets of neurons by targeting neuropeptide loci

We determined if this CRISPR/Cas9-mediated insertion could be used to build killifish reporter lines for specific cell types, notably neuronal subpopulations. This development is critical for systems neuroscience, including circuit-based studies. We focused on targeting neurons expressing neuropeptide Y (NPY) and hypocretin (HCRT). These neuronal populations are critical for organismal homeostasis through modulation of behaviors, including feeding behavior (Jeong et al., 2018) and sleep-wake behavior (Chiu and Prober, 2013; Prober et al., 2006; Singh et al., 2017). Growing evidence also suggests that these neuronal populations may be altered with age (Fronczek et al., 2012; Hunt et al., 2015; Montesano et al., 2019). We designed a dsDNA HDR template encoding Venus targeting the *NPY* or *HCRT* locus and with a *T2A* sequence 5’ to the fluorescent protein sequence to avoid direct fusion between the neuropeptide and the fluorescent protein (Figure 4A; Supplemental Table 1). PCR amplification and genotyping by Sanger sequencing of F1 animals confirmed that the *T2A*-*Venus* integration at the *NPY* or *HCRT* locus was as designed—single-copy and in frame at the targeted genomic location (Figure 4A; Figure 4—figure supplement 1; Supplemental Table 1). Imaging of coronal brain sections of the adult *HCRT*-*T2A*-*Venus* killifish line showed a dense and isolated population of Venus-positive cell bodies in the dorsal periventricular hypothalamus (Hd; homologous to the mammalian arcuate nucleus) (Appelbaum et al., 2009; Biran et al., 2015; D’Angelo, 2013; Montesano et al., 2019) (Figure 4B). In contrast, the adult *NPY*-*T2A*-*Venus* line exhibited Venus-positive cell bodies throughout the brain, including in the periventricular and lateral hypothalamus, as well as in the periventricular gray zone (PGZ) of the optic tectum (OT) (Figure 4C). The expression profiles observed in the *NPY*-*T2A*-*Venus* and *HCRT*-*T2A*-*Venus* lines are consistent with *in situ* hybridization of endogenous *NPY* and *HCRT* transcripts in wildtype animals (Figure 4—figure supplement 2; Supplemental Table 3), and also consistent with *NPY* and *HCRT* expression previously reported in the adult killifish and zebrafish brain (Appelbaum et al., 2009; Biran et al., 2015; D’Angelo, 2013; Montesano et al., 2019). The generation of these lines serves as proof of principle that CRISPR/Cas9-mediated knock-in is a powerful method in killifish to drive cell-type-specific expression. These neuron-specific lines should also help the development of the killifish for systems neuroscience studies.

## Discussion

Here we establish an efficient and versatile method for rapid and precise genome engineering of the short-lived African turquoise killifish. This CRISPR/Cas9-mediated knock-in method can be leveraged for cell-type-and tissue-specific expression of ectopic genes and reporters to study complex phenotypes at scale. We observe efficient CRISPR/Cas9-mediated knock-in of large inserts (>40% efficiency) with germline transmission rates over 65%. This high efficiency of germline transmission may be due to the relatively slow rate of early cell division after fertilization in the African turquoise killifish (∼4 times slower in this species relative to non-annual teleost fishes) (Dolfi et al., 2014). The killifish model, with hundreds of embryos produced at a given time (for example using harem breeding), allows for easy and high-throughput injection of genome-editing machinery into embryos (Harel et al., 2015; Hu and Brunet, 2018; Kim et al., 2016; Polacik et al., 2016). Moreover, the killifish has the shortest generation time of any vertebrate model bred in captivity (Hu and Brunet, 2018; Kim et al., 2016). The development of rapid and efficient knock-in establishes the killifish as a system for precise genetic engineering at scale, which has been challenging so far in vertebrates. The knock-in method developed here uses reagents that are all commercially available, eliminating the need for cloning and PCR and making this method easy to adopt. Together, the steps described here could serve as a blueprint for knock-in approaches in other emerging model organisms.

The CRISPR/Cas9-mediated knock-in approach we developed should allow the establishment of versatile strategies to probe complex phenotypes, including development, ‘suspended animation’, regeneration, aging, and age-related diseases. Given the potential of the African killifish for modeling human aging (Hu and Brunet, 2018; Kim et al., 2016; Van Houcke et al., 2021a), this knock-in method should also allow the generation of human disease models that can be studied longitudinally, over an entire lifespan. For example, this CRISPR/Cas9-mediated knock-in could be used to introduce human neurodegenerative disease variants into conserved endogenous killifish loci (e.g., amyloid precursor protein [*APP*] for Alzheimer’s disease) or to drive neurodegenerative disease variants using a pan-neuronal promoter. Human disease variant models in mice have been critical to understand disease mechanisms and treatment strategies (Dawson et al., 2018; Fisher and Bannerman, 2019; Jankowsky and Zheng, 2017). Human disease models that are scalable and integrate both genetics and age as risk factors have the potential to identify new strategies to treat these diseases.

This study highlights the power of knock-in, combined with self-cleaving peptides, to drive cell-type-specific expression of ectopic genes such as molecular reporters (e.g., fluorescent reporters, calcium indicators), recombinases (e.g., Cre), and optogenetic tools (e.g., light sensitive ion channels such as channelrhodopsin). The cell-type resolution of this genetic tool should open studies in a variety of fields, including systems neuroscience. Additional variations, such as the use of ‘landing pads’ (for higher levels of expression) (Soriano, 1999) and inducible promoters (either endogenous or ectopic) (Gossen and Bujard, 1992; Gossen et al., 1995), could be further developed to complete this toolkit. Overall, this knock-in method should accelerate the use of the killifish as a scalable vertebrate model and allow discoveries in several fields, including regeneration, neuroscience, aging, and disease, with conserved implications for humans.

## Supplemental Figures

**Figure 1—figure supplement 1:**
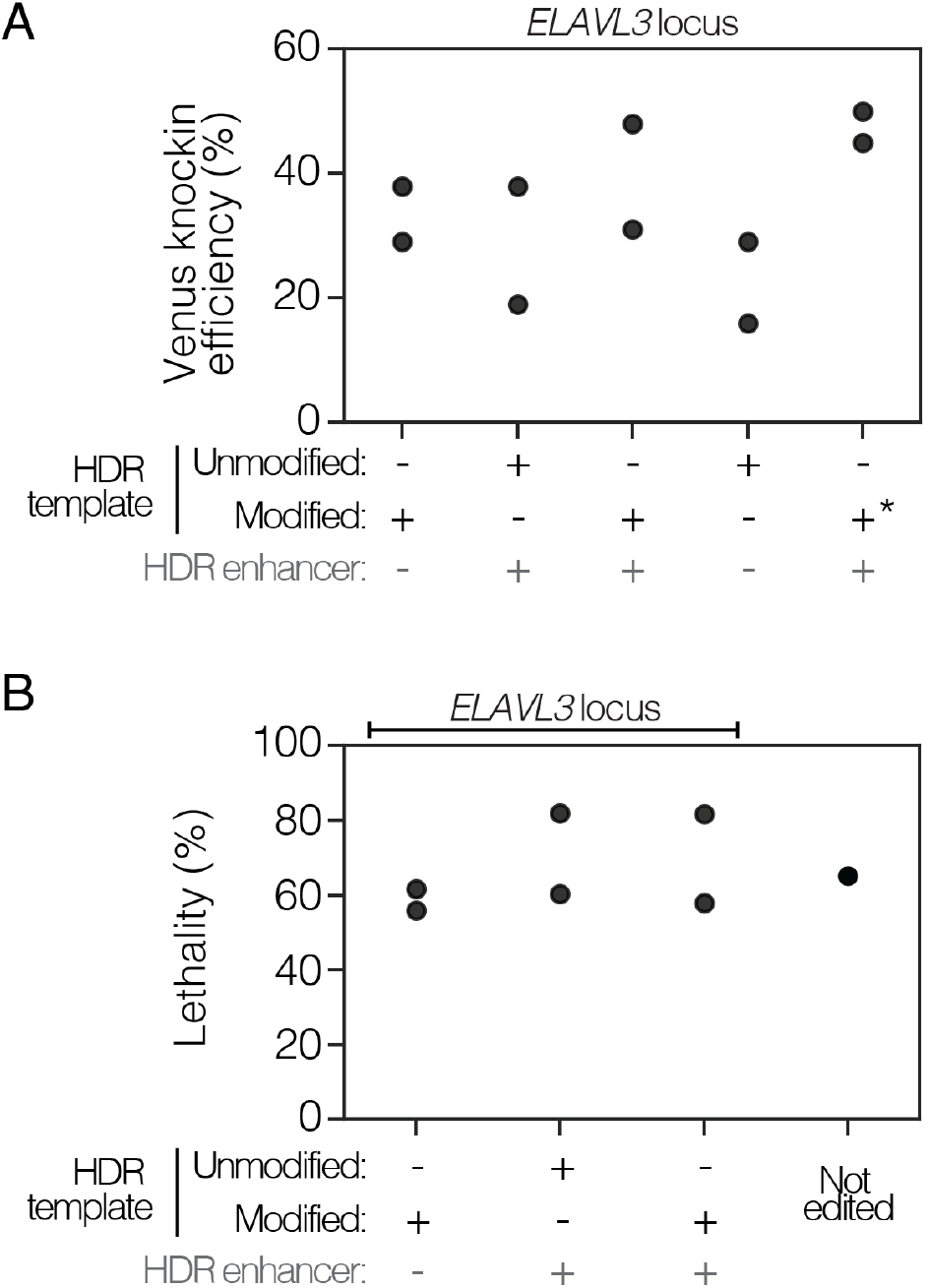
Comparing knock-in efficiency and lethality of chemically modified dsDNA HDR templates and HDR chemical enhancers. A. Knock-in efficiency of different knock-in reagents for insertion of *T2A*-*Venus* at the *ELAVL3* locus comparing the use of chemically modified dsDNA HDR templates versus unmodified dsDNA HDR templates, and the use of HDR chemical enhancers (+/-HDR enhancer). Two types of HDR modifications were tested: modification #1 and modification #3. The use of modification #3 is indicated by (*). Modification #3 was used for all subsequent work in this paper (see Methods). Two replicates per condition; *n* = 10–105 embryos per replicate. Raw data in Supplemental Table 2. B. Lethality over two weeks after CRISPR/Cas9-mediated knock-in with different knock-in reagents for insertion of *T2A*-*Venus* at the *ELAVL3* locus, compared to non-injected (not edited) control. Two replicates per knock-in condition. Raw data in Supplemental Table 2.

**Figure 2—figure supplement 1:**
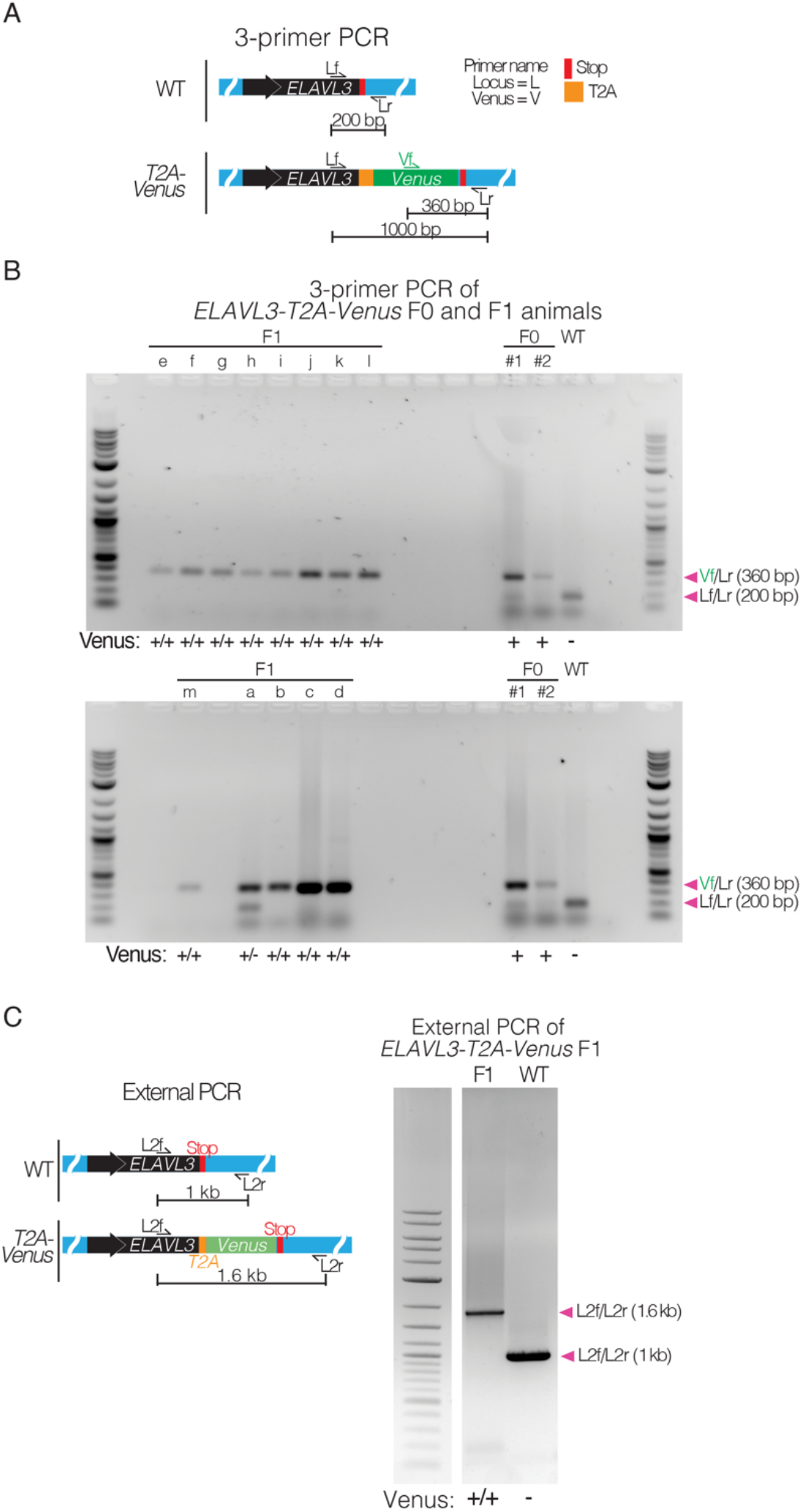
PCR amplification of F0 parents and F1 progeny confirms *T2A-Venus* integration and germline transmission at the *ELAVL3* locus. A. 3-primer PCR schematic showing locus-specific external primers forward (Lf) and reverse (Lr) and internal Venus forward primer (Vf) at the *ELAVL3* locus. B. Crossing F0 animals positive for *T2A*-*Venus* at the *ELAVL3* locus (F0 X F0). Gels showing F0 parents (right) and F1 progeny (a, b, c, and d; left) with 3-primer PCR using Venus and locus-specific primers shown in (A). Arrowheads indicate each primer pair and its expected amplification product length. Scoring for each lane of the gel is indicated below the gel images. Full gel images of 3-primer PCR on F0 and F1 animals for gels shown in Figure 2C. C. PCR amplification at the *ELAVL3* locus of F1 homozygous *ELAVL3*-*T2A*-*Venus* killifish compared to WT with forward and reverse primers external to the homology arms (i.e., external PCR). *ELAVL3*-*T2A*-*Venus* killifish produce a band at the expected length (1.6 kb). Arrowheads indicate each primer pair and its expected amplification product length. Scoring for each well of the gel is indicated below the gel images.

**Figure 2—figure supplement 2:**
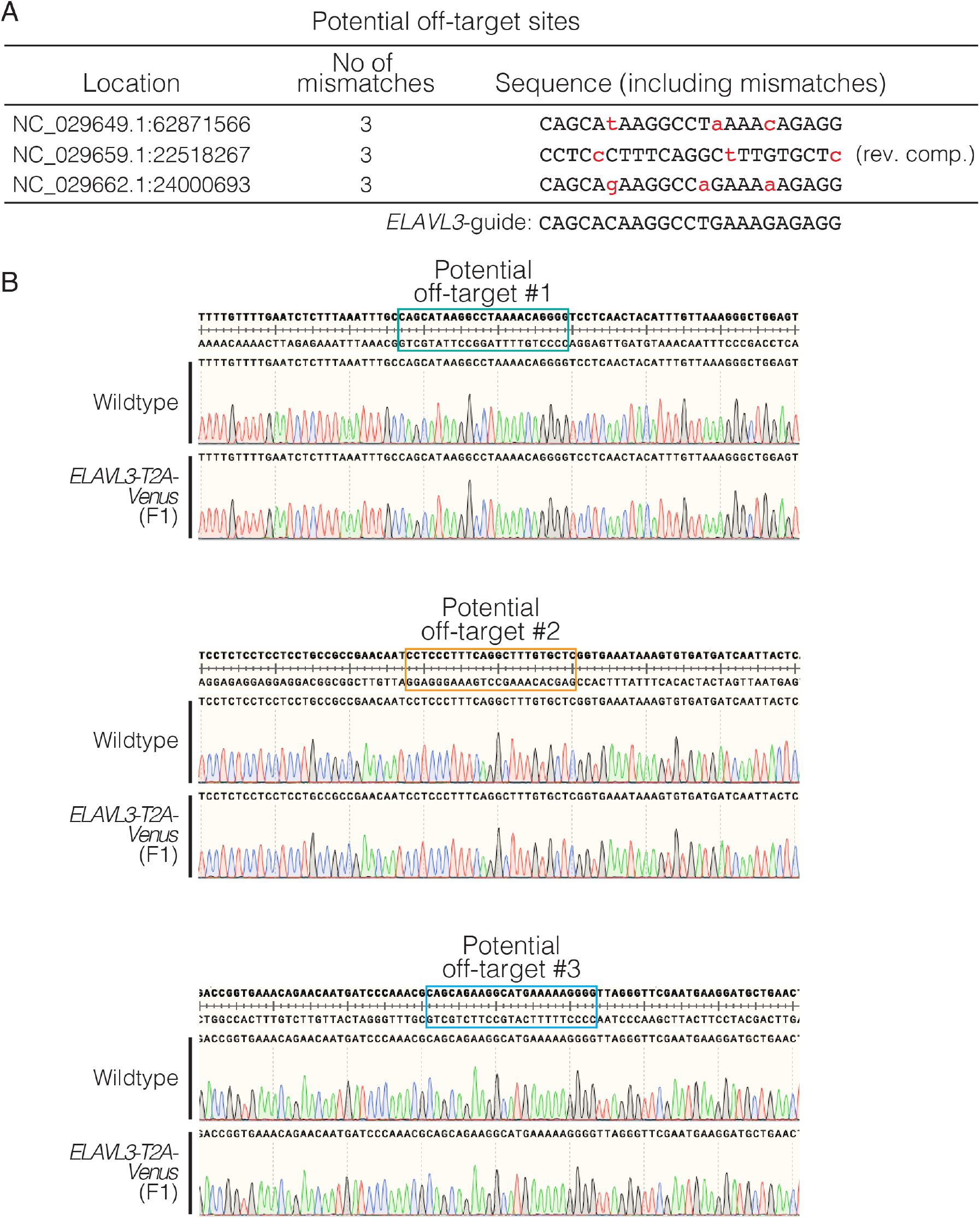
Evaluating potential off-target effects in homozygous F1 CRISPR/Cas9 knock-in fish. A. The predicted most likely off-target sites for the *ELAVL3* gRNA (predicted using CHOPCHOP). These three loci each have three mismatches from the *ELAVL3* gRNA (indicated by lowercase, red text). B. Sequencing results of *ELAVL3-T2A-Venus* homozygous F1 (generated by crossing F0s) at the three predicted most likely off-target sites. No off-target editing was observed at any of the predicted off-target sites.

**Figure 3—figure supplement 1:**
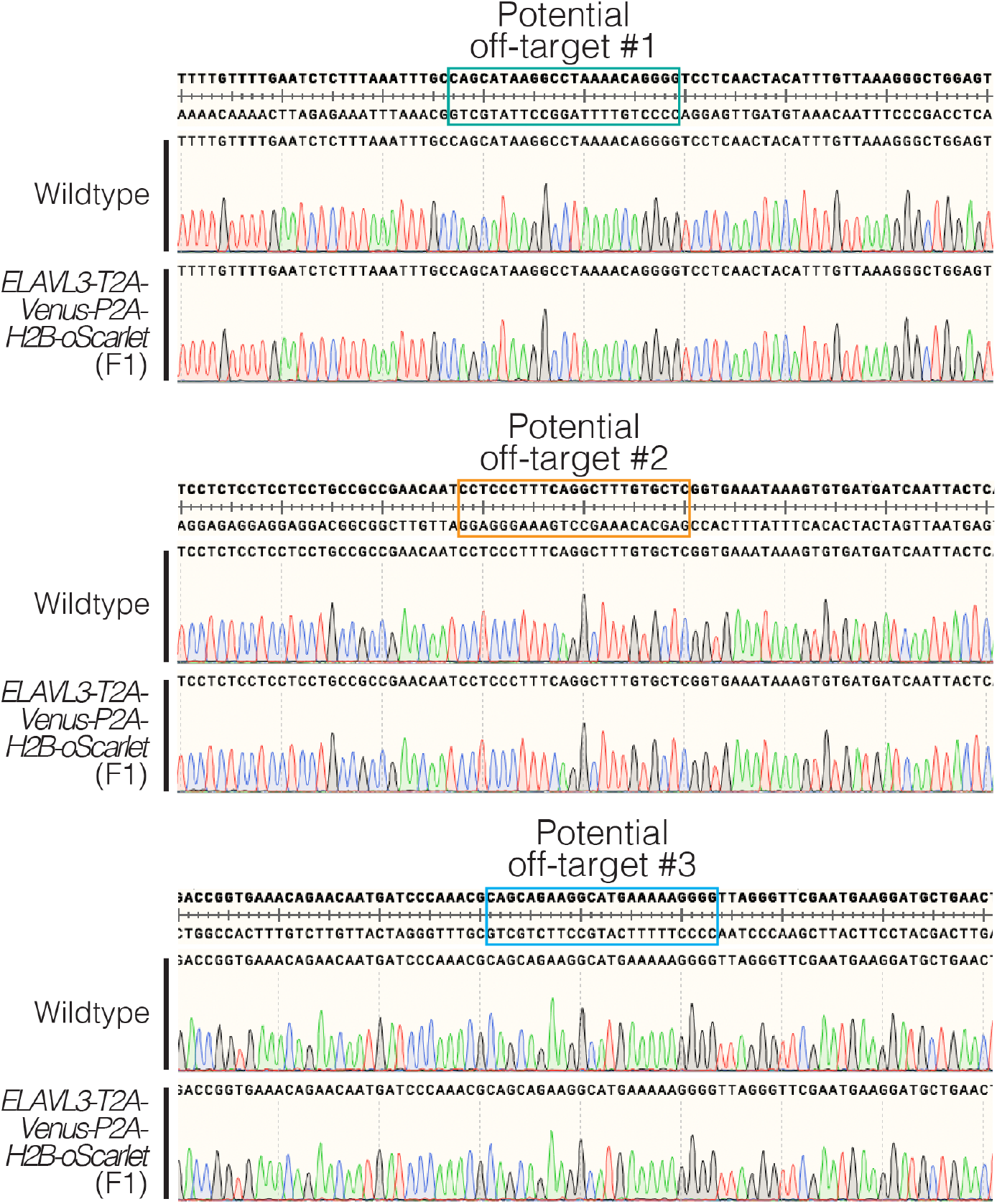
Evaluating potential off-target effects in homozygous F1 CRISPR/Cas9 knock-in fish. Sequencing results of *ELAVL3-T2A-Venus-P2A-H2B-oScarlet* homozygous F1 (generated by crossing F0s) at the three predicted most likely off-target sites. No off-target editing was observed at any of the predicted off-target sites.

**Figure 4—figure supplement 1:**
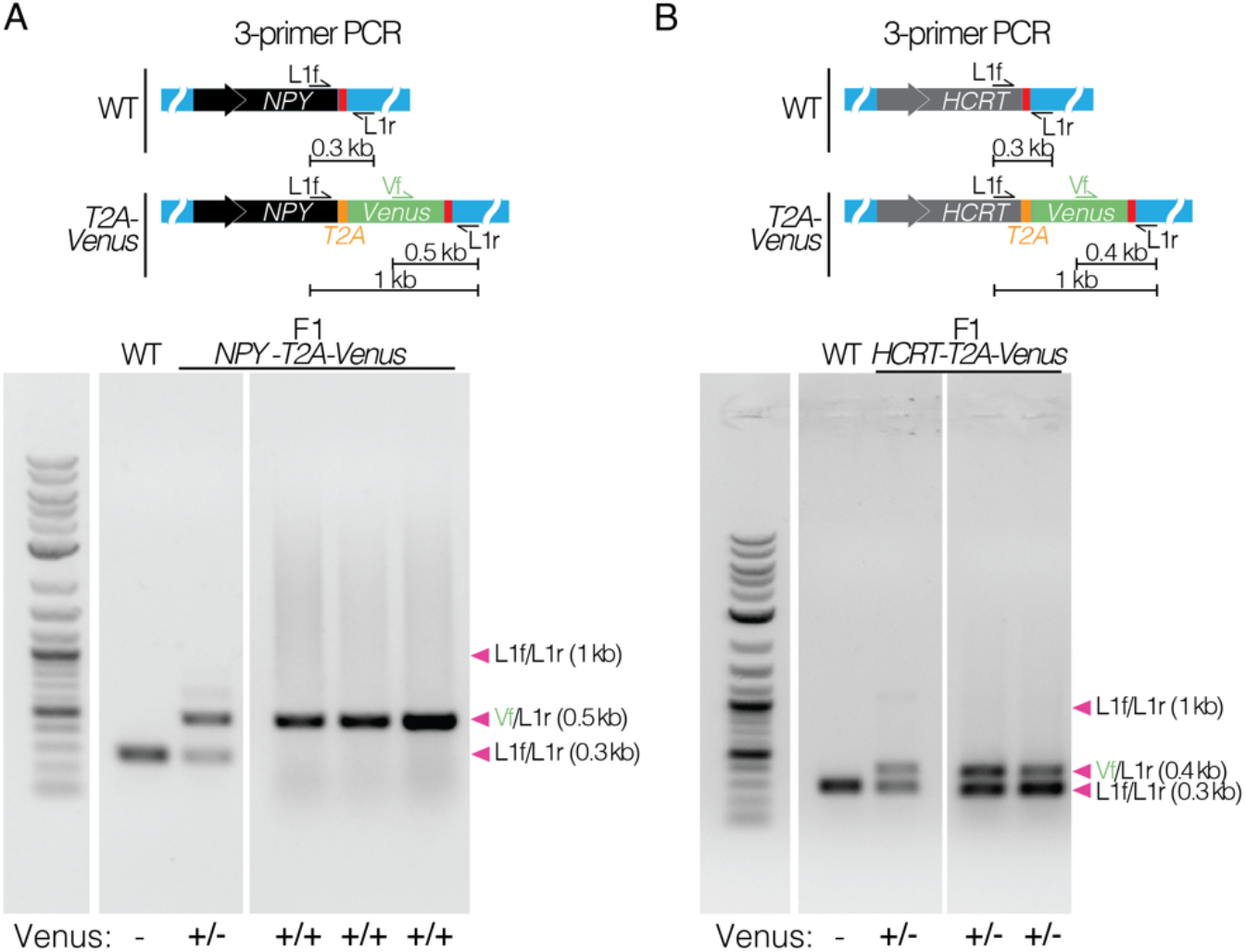
Confirming knock-in by PCR amplification and sequencing. A. 3-primer PCR amplification at the *NPY* locus of F1 heterozygous and homozygous *NPY*-*T2A*-*Venus* killifish compared to WT. Arrowheads indicate each primer pair and its expected amplification product length. Scoring Venus negative (-), heterozygous (+/-), or homozygous (+/+) for each well of the gel is indicated below the gel images. The PCR product was sequenced and confirmed. B. 3-primer PCR amplification (schematic and gel images) at the *HCRT* locus of F1 heterozygous *HCRT*-*T2A*-*Venus* killifish compared to WT. Arrowheads indicate each primer pair and its expected amplification product length. Scoring Venus negative (-), heterozygous (+/-), or homozygous (+/+) for each well of the gel is indicated below the gel images. The PCR product was sequenced and confirmed.

**Figure 4—figure supplement 2:**
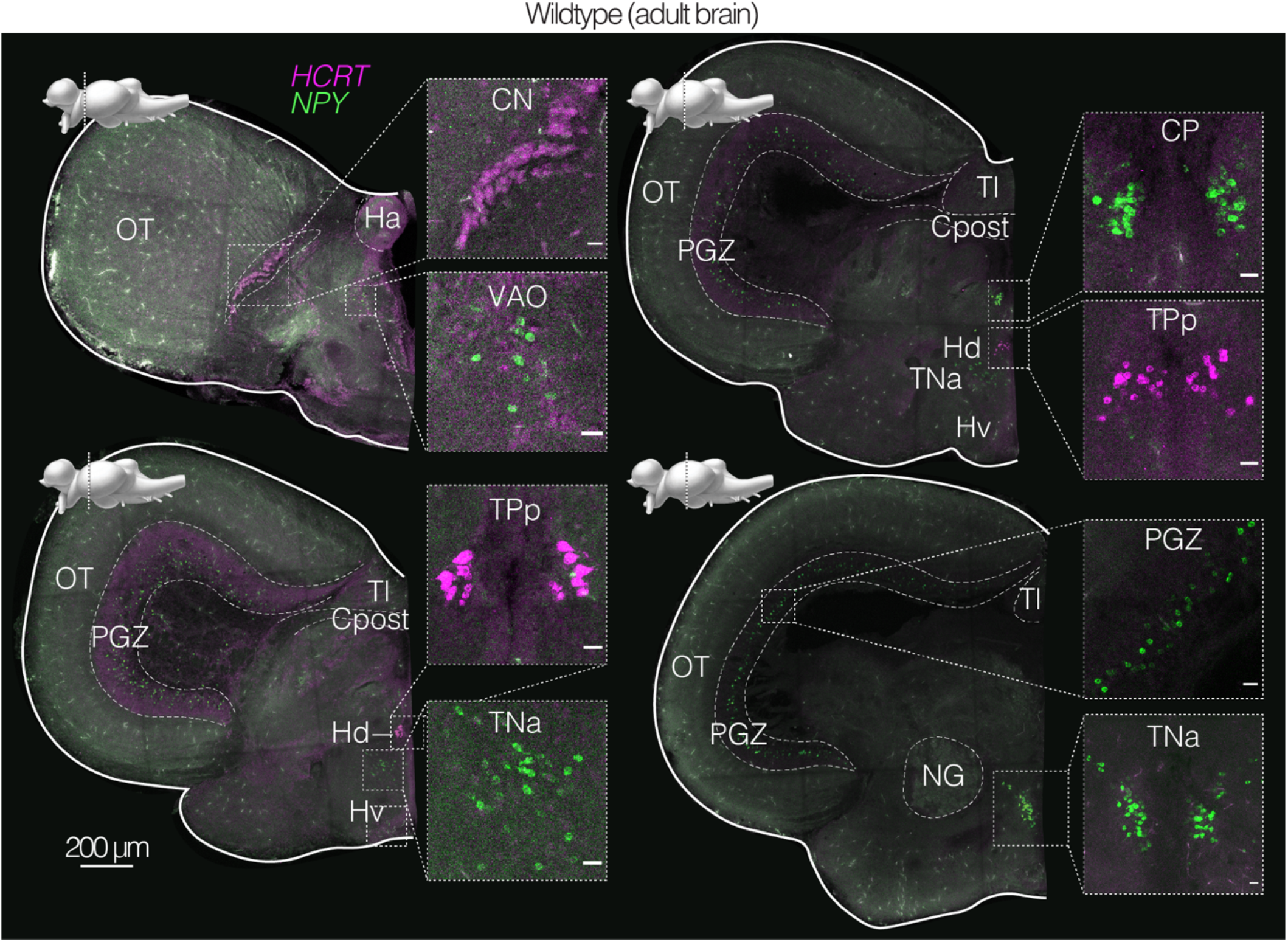
*In situ* hybridization chain reaction (HCR) of endogenous *NPY* and *HCRT* transcripts in the adult brain of wildtype killifish. Coronal brain sections of an adult (3 months old) wildtype male, with HCR labeling endogenous *NPY* (green) and *HCRT* (magenta) transcripts (using gene-specific probe sets, Supplemental Table 3). Scale bar = 200 μm. Distinct nuclei indicated and labeled with abbreviated names. Above each slice is the sagittal view of the *N. furzeri* brain adapted from (D’Angelo, 2013) indicating the plane of the coronal section. Inset shows zoom in on *NPY* and *HCRT* positive population of cells. Scale bar = 20 μm.

## Methods

### African turquoise killifish care and husbandry

African turquoise killifish (GRZ strain) were maintained according to established guidelines (Astre et al., 2022b; Reichard et al., 2022; Zak et al., 2020). Briefly, animals were housed at 26–27°C in a central filtration recirculating system (Aquaneering, San Diego) at a conductivity between 3800–4000 μS/cm and a pH between 6.5–7.0, with a daily exchange of 10% water treated by reverse osmosis (i.e., RO water). Animals were kept on a 12-hour light/dark cycle and were fed twice a day on weekdays and once a day on weekends. Adult fish (>1 month of age) were fed dry fish food (Otohime fish diet, Reed Mariculture, Otohime C1) while young fish (<1 month of age) were fed freshly hatched brine shrimp (Brine Shrimp Direct, BSEP6LB). Killifish embryos were raised in Ringer’s solution (Sigma-Aldrich, 96724), with two tablets per liter of RO water and 0.01% methylene blue (i.e., embryo solution) at 26–27°C in 60 mm x 15 mm petri dishes (E and K Scientific, EK-36161) at a density of <100 embryos per plate. After two weeks in embryo solution, embryos were transferred to moist autoclaved coconut fiber (Zoo Med Eco Earth Loose Coconut Fiber) lightly packed in petri dishes where they were incubated for another two weeks at 26–27°C. After 2–3 weeks on moist coconut fiber, embryos were hatched. For hatching, embryos were placed in humic acid solution (1 g/l, Sigma-Aldrich, 53680 in RO water) and incubated overnight at room temperature. All animals were raised in accordance with protocols approved by the Stanford Administrative Panel on Laboratory Animal Care (protocol # APLAC-13645).

### Design of guide RNA sequences

For each selected gene, gRNA target sites were identified using CHOPCHOP (Labun et al., 2019) (https://chopchop.rc.fas.harvard.edu/) with the Nfu_20140520/Jena genome. One guide sequence was selected for each target gene of interest. Guide sequences were only selected if followed by the PAM site (5’-NGG-3’) for *Streptococcus pyogenes* Cas9. The Cas9 cut sites were between 1–15 bp from the target insertion site. Guide RNAs were designed for compatibility with Integrated DNA Technology’s (IDT, Coralville, IA) Alt-R™ method. For detailed methods and design tools see www.idtdna.com. All Alt-R CRISPR RNAs (crRNAs) and universal trans-activating crRNA (tracrRNA) were chemically synthesized (2 nmol, IDT). Synthetic Alt-R™ crRNA and tracrRNA were resuspended in nuclease-free duplex buffer (IDT) to a final concentration of 100 μM each and stored at -20°C. The guide sequences of all crRNAs used in this study are provided in Supplemental Table 1.

### Design of DNA templates for HDR

Double-stranded DNA (dsDNA) HDR templates were designed with 150–200 bp homology arms containing DNA sequences surrounding the target Cas9 cut site. Homology arms began within 1–15 bp of the Cas9 cut site. All dsDNA HDR templates were synthesized as gBlock Gene Fragments from IDT (0.25–10 μg). Unless otherwise noted, all gBlocks contained IDT’s proprietary chemical modifications at each end of the sequence that should promote HDR and inhibit blunt-end integration. In this work, we tested two proprietary chemical modifications from IDT. We found chemical modification #3 worked best and used it for the majority of the HDR templates used in this study (Figure 1—figure supplement 1A). gBlocks with this modification #3 are available for purchase through IDT as Alt-R™ HDR Donor Blocks. gBlocks were resuspended in nuclease-free duplex buffer (IDT) to a concentration of 150 ng/μl and stored at -20°C. HDR template sequences used in this study are provided in Supplemental Table 1.

### Preparation and microinjection of CRISPR/Cas9 reagents into African turquoise killifish embryos

To prepare the gRNA complex, the tracrRNA and crRNA were mixed in a 1:1 ratio for a final concentration of 3 μM in nuclease free duplex buffer and annealed by incubation at 95°C for 5 minutes followed by cooling to room temperature. To form the ribonucleoprotein (RNP) complex, the gRNA complex was mixed with Cas9 protein (IDT, 1081059; 10 μg/μl) in 1x phosphate-buffered saline (1xPBS; Corning, 21-040-CV) to final concentrations of 1.5 μM gRNA complex and 0.25 μg/μl Cas9 protein. This mixture was then incubated at 37°C for 10 minutes followed by cooling to room temperature. The chemically-modified dsDNA HDR template and IDT’s HDR chemical enhancer (IDT’s Alt-R HDR enhancer Version 2 [V2]) were then added to the RNP complex for injection, for final concentrations of 50 ng/μl gRNA complex, 250 ng/μl Cas9 protein, and 15 ng/μl HDR template. Finally, 0.33 μl of 8% phenol red was added to the injection mixture for visualization. The mixture was used immediately (within 1 hour of production) and kept on ice. Preassembled Cas9 RNP complex and synthetic dsDNA HDR templates were injected into the single cell of one-cell stage killifish embryos in accordance with microinjection procedures described in (Harel et al., 2015). For each target locus and HDR template, 60–150 embryos were injected. Surviving injected embryos were maintained in embryo solution at 26–27°C for 2–3 weeks. Embryos were then transferred to moist autoclaved coconut fiber (Zoo Med Eco Earth Loose Coconut Fiber) lightly packed in petri dishes where they were incubated for another 2–3 weeks at 26–27°C after which they were hatched (as described in <u>African turquoise killifish care and husbandry</u>).

### Assessment of genome editing

Visual screening: Visual fluorescence screening of 14–21-day old F0 embryos on a Fluorescent Stereo Microscope (Leica M165FC; Figure 1B) was conducted to verify successful knock-in of cDNA encoding fluorescent proteins. Twenty-one-day old embryos were dried on coconut fiber for seven days prior to imaging.

Genotyping: PCR amplification of genomic DNA from fish tail clips was also used to verify successful knock-in events. For this, we followed protocol described in (Hu et al., 2020). Briefly, caudal fin clips were taken from 1–3-week-old fish (anesthetized on ice). Clipped fin tissue was digested in 30 μl DirectPCR Lysis Reagent (Mouse Tail) (Viagen, 102-T) with 40 μg/ml Proteinase K (Invitrogen, 25530049) at 55°C for 2 hrs followed by 100 °C heat inactivation for 10 min. This solution was used as template for PCR amplification with the following PCR reaction mixture (20 μl): 3 μl crude tail-clip lysis, 1 μl 100 μM primers (IDT), 10 μl 2x GoTaq® Master Mixes (Promega, M7123), and 6 μl water. The PCR was run for 30–42 cycles. We used primer sets that enabled detection of genome editing based on amplification product size by gel electrophoresis (Figure 1D; Figure 2C; Figure 2—figure supplement 1; Figure 3C; Figure 4A; Figure 4—figure supplement 1). The primer sequences used to verify successful editing by genotyping are provided in Supplemental Table 1.

Sequencing: To verify the sequence of successfully edited genomes, PCR amplification of the genomic DNA from fish tail clips was also sent for sequencing (Molecular Cloning Laboratories, MCLAB, https://www.mclab.com/home.php). The sequencing primer sequences used to verify successful editing are provided in Supplemental Table 1.

### Tissue histology

For brain sectioning and staining, extracted whole brain samples from 1–4-month-old animals were fixed overnight in 4% paraformaldehyde in PBS (Santa Cruz Biotechnology, SC281692) at 4°C and then washed for 12 hrs in 1xPBS (Corning, 21-040-CV) at 4°C with three washes. Fixed samples were dehydrated in 30% sucrose (Sigma-Aldrich, S3929) in 1xPBS at 4°C overnight or until tissue sunk. Tissue was then embedded in Tissue-Plus™ OCT (Fisher Scientific, 23-730-571) within plastic embedding molds. Tissue was then frozen at -20 °C for at least 2 hrs and sectioned (50–100 μm sections) on a cryostat (Leica CM3050 S) and mounted on glass slides (Fisher Scientific, 12-550-15) and stored at - 20°C.

For immunostaining, slides were washed once in 1xPBS at room temperature to remove residual OCT. Slides were dehydrated and permeabilized in pre-chilled 100% methanol (Sigma-Aldrich, HPLC grade) with 1% Triton X-100 (Fisher Scientific, BP151) at -20°C for 15 min, followed by washing in 1xPBS at room temperature. Slides were blocked with 5% Normal Donkey Serum (NDS; ImmunoReagents Inc., SP-072-VX10) and 1% Bovine Serum Albumin (BSA; Sigma, A7979) in 1xPBS (“blocking buffer”) for 30 min at room temperature. Slides were washed in 1xPBS with 0.1% Tween-20 (PBST) three times for 10 min each, followed by washing in PBS. Slides were incubated in primary antibody (rabbit GFP Polyclonal Antibody, ThermoFisher, A-6455) at a 1:250 dilution in blocking buffer overnight at 4°C followed by washing in PBST three times for 30 min each and then washing in 1xPBS. Slides were incubated in secondary antibody (donkey anti-rabbit IgG, ThermoFisher, A-31573) at a 1:500 dilution in blocking buffer for 2 hrs at room temperature followed by washing in PBST three times for 30 min each and then washing in 1xPBS. Slices were mounted in either ProLong™ Gold Antifade Mountant (ThermoFisher, P36930) or ProLong Gold Antifade Mountant with DAPI (ThermoFisher, P36931) for imaging.

### *In situ* hybridization chain reaction (HCR)

For in situ hybridization by hybridization chain reaction (HCR), we followed a protocol described in (Lovett-Barron et al., 2017). First, hybridization probes were designed according to the split initiator approach of third generation in situ hybridization chain reaction (Choi et al., 2018), which enables automatic background suppression. Twenty-two-nucleotide long DNA antisense oligonucleotide split probes were designed for both *NPY* and *HCRT* based on the killifish mRNA sequence (Supplemental Table 3) and synthesized by IDT (200 μM in RNAse-free H_2_O). Dye-conjugated hairpins (B3-488 and B5-546) were purchased from Molecular Instruments. Slides were washed in 1xPBS at room temperature to remove residual OCT. Slides were dehydrated and permeabilized in pre-chilled 100% methanol (Sigma-Aldrich, HPLC grade) with 1% Triton X-100 (Fisher Scientific, BP151) at -20°C for 15 min followed by washing three times in 2X saline sodium citrate (SSC) buffer with 0.1% Tween-20 (2xSSCT; made from 20xSSC, ThermoFisher, AM9763) at room temperature for 30 min each.

Slides were equilibrated in hybridization buffer (2xSSCT, 10% (w/v) dextran sulfate [Sigma Aldrich, D6001], 10% (v/v) formamide [Thermo Fisher, AM9342]) for 30 min at 37°C. Slides were then hybridized with split probes in hybridization buffer at a probe concentration of 4 nM overnight at 37°C. Slices were then washed two times in 2xSSCT and 30% (v/v) formamide for 30 min at 37 °C. Slides were washed two times in 2xSSCT for 30 min each at room temperature. Slides were pre-amplified in amplification buffer (Molecular Instruments) for 10 min at room temperature. Dye-conjugated hairpins were prepared according to manufacturer’s instructions. Briefly, they were heated to 95°C for 1 min then snap-cooled to 4°C. Amplification was performed by incubating slides in amplification buffer with prepared B3 and B5 probes at concentrations of 120 nM overnight in the dark at room temperature. Slides were washed 3 times with 2xSSCT for 30 min each. Slices were mounted in ProLong™ Gold Antifade Mountant for imaging.

### Whole-mount tissue clearing

For whole-mount tissue clearing (shown in Figure 3E), extracted whole brain samples from 1–4-month-old animals were fixed overnight in 4% paraformaldehyde in 1xPBS at 4°C and then washed for 12 hrs in 1xPBS 4°C with three washes. Fixed brain samples were crosslinked in a SHIELD hydrogel (Park et al., 2018) overnight in 1–2% SHIELD epoxide reagent (GE38; CVC Thermoset Specialties of Emerald Performance Materials) in 0.1 M Carbonate Buffer (pH 8.3) at 37°C and then washed three times for 1 h each in 1xPBS at 37°C. Samples were cleared for 12–48 hrs (depending on brain size) in 4% sodium dodecyl sulfate (SDS) at 37°C until optically translucent and then washed three times for 1 h intervals in 1xPBS with 0.1% Tween-20 (PBST) at 37°C. For imaging, samples were then equilibrated in EasyIndex (RI = 1.52, LifeCanvas Technologies) and mounted.

### Imaging

All samples (unless otherwise noted) were imaged using an Olympus FV1200 confocal microscope system running Fluoview software, using a 10x 0.6 Numerical Aperture water immersion Olympus objective. Images were collected at a 5 μm z-step resolution. For higher magnification images in Figure 3D, samples were imaged using a Zeiss LSM900 confocal microscope (Axio Observer) system running ZEN software (3.0, blue), using a 40x 1.4 Numerical Aperture oil immersion Zeiss objective (Plan-Apochromat). Images were collected at a 4.5 μm z-step resolution. Single photon excitation was used at the indicated wavelengths. Entire samples were obtained by mosaic tiling during imaging, reconstructed using Fluoview software, and viewed and analyzed in Fiji and Aivia software.

## Abbreviations

CN: cortical nucleus
CP: central posterior thalamic nucleus
Cpost: posterior commissure
DIL: diffuse inferior lobe of hypothalamus
gl: glomerular layer
Ha: habenular nucleus
Hc: caudal hypothalamus
Hd: dorsal hypothalamus
Hv: ventral hypothalamus
llf: lateral longitudinal fascicle
LR: lateral recess of diencephalic ventricle
mlf: medial longitudinal fascicle
MO: medulla oblongata
NG: glomerular nucleus
OB: olfactory bulb
ON: optic nerve
OT: optic tectum
PGZ: periglomerular gray zone
Tel: telencephalon
Tl: torus longitudinalis
TNa: anterior tuberal nucleus
TPp: periventricular nucleus of posterior tuberculum
Va: valvula of cerebellum
VAO: ventral accessory optic nucleus

## Acknowledgements

We thank Drs. Felix Boos, Jing Chen, Tyson Ruetz, and John Bedbrook, as well as Lucy Xu and Charu Ramakrishnan, and all members of the Brunet lab and Deisseroth lab for their input on the project and providing feedback on the manuscript. We thank IDT for generously providing reagents for testing. We thank Rogelio Barajas, Rishad Khondker, Jacob Chung, and Natalie Schmahl for killifish husbandry support. We thank Rogelio Barajas for managing the killifish room and help with development and maintenance of lines. This work was supported by R01AG063418 (A.B. and K.D.), the Glenn Foundation for Medical Research (A.B.), the Simons Foundation (A.B.), the Knight-Hennessy Scholars Graduate Fellowship (R.N.), the Helen Hay Whitney Postdoctoral Fellowship (C.N.B), the Wu Tsai Stanford Neuroscience Institute Interdisciplinary Postdoctoral Fellowship (C.N.B), T32 AG0047126 (R.D.N.), and the Iqbal Farrukh & Asad Jamal Center for Cognitive Health in Aging (R.D.N).

## Author contributions

R.D.N. and C.N.B. conceptualized the project, with input from K.D. and A.B.. R.D.N. and C.N.B. designed, performed, and analyzed all experiments with help from R.N.. R.D.N. designed gRNAs and DNA templates for HDR, performed injections for generating lines, and managed and maintained lines with assistance from C.N.B. and R.N.. C.N.B. developed the protocol for CRISPR/Cas9-mediated knock-in and performed tissue histology and imaging. R.N. optimized and performed genotyping and quantified knock-in efficiency with help and guidance from C.N.B and R.D.N..R.D.N. and C.N.B. wrote the manuscript with help from R.N. and A.B.. All the authors provided intellectual input and commented on the manuscript.

